# Opposing activities of IFITM proteins in SARS-CoV-2 infection

**DOI:** 10.1101/2020.08.11.246678

**Authors:** Guoli Shi, Adam D. Kenney, Elena Kudryashova, Lizhi Zhang, Luanne Hall-Stoodley, Richard T. Robinson, Dmitri S. Kudryashov, Alex A. Compton, Jacob S. Yount

**Affiliations:** HIV Dynamics and Replication Program, National Cancer Institute, Frederick, MD, USA; Department of Microbial Infection and Immunity, The Ohio State University College of Medicine, Columbus, OH, USA; Viruses and Emerging Pathogens Program, Infectious Diseases Institute, The Ohio State University, Columbus, OH, USA; Department of Chemistry and Biochemistry, The Ohio State University, Columbus, OH, USA

## Abstract

Interferon-induced transmembrane proteins (IFITMs) restrict infections by many viruses, but a subset of IFITMs enhance infections by specific coronaviruses through currently unknown mechanisms. Here we show that SARS-CoV-2 Spike-pseudotyped virus and genuine SARS-CoV-2 infections are generally restricted by expression of human IFITM1, IFITM2, and IFITM3, using both gain- and loss-of-function approaches. Mechanistically, restriction of SARS-CoV-2 occurred independently of IFITM3 *S*-palmitoylation sites, indicating a restrictive capacity that is distinct from reported inhibition of other viruses. In contrast, the IFITM3 amphipathic helix and its amphipathic properties were required for virus restriction. Mutation of residues within the human IFITM3 endocytosis-promoting YxxΦ motif converted human IFITM3 into an enhancer of SARS-CoV-2 infection, and cell-to-cell fusion assays confirmed the ability of endocytic mutants to enhance Spike-mediated fusion with the plasma membrane. Overexpression of TMPRSS2, which reportedly increases plasma membrane fusion versus endosome fusion of SARS-CoV-2, attenuated IFITM3 restriction and converted amphipathic helix mutants into strong enhancers of infection. In sum, these data uncover new pro- and anti-viral mechanisms of IFITM3, with clear distinctions drawn between enhancement of viral infection at the plasma membrane and amphipathicity-based mechanisms used for endosomal virus restriction. Indeed, the net effect of IFITM3 on SARS-CoV-2 infections may be a result of these opposing activities, suggesting that shifts in the balance of these activities could be coopted by viruses to escape this important first line innate defense mechanism.

## Introduction

Severe acute respiratory syndrome (SARS) coronavirus (CoV)-2 is the causative agent of the respiratory and multi-organ-associated disease known as COVID-19 (Wu, Zhao et al., 2020, Zhu, Zhang et al., 2020). Understanding virus-host interactions and immune responses will be critical for explaining the high variation in severity of COVID-19 observed in humans (Zhang, Tan et al., 2020). Type I interferon (IFN) is a critical component of the innate immune system that induces expression of hundreds of genes, many of which encode proteins with antiviral activities that block specific steps in viral replication cycles (Chemudupati, Kenney et al., 2019, Schoggins, Wilson et al., 2011). Among these are genes that encode the IFN-induced transmembrane proteins (IFITMs), including IFITM1, IFITM2, and IFITM3. IFITMs have been shown to block membrane fusion of diverse enveloped viruses by using an amphipathic helix domain to mechanically alter membrane lipid order and curvature in a manner that disfavors virus fusion (Chesarino, Compton et al., 2017, Guo, Steinkühler et al., 2020, Li, Markosyan et al., 2013, Lin, Chin et al., 2013). While IFITMs have been shown in most instances to inhibit CoV infections (Bertram, Dijkman et al., 2013, Huang, Bailey et al., 2011, Wrensch, Winkler et al., 2014, Zheng, Zhao et al., 2020), in some cases they enhance infections(Zhao, Guo et al., 2014, Zhao, Sehgal et al., 2018). For example, IFITM2 and IFITM3 enhance cellular entry of human CoV-OC43 (Zhao et al., 2014). Similarly, infections by virus-like particles containing SARS-CoV or MERS-CoV glycoprotein (pseudovirus) are increased upon expression of an IFITM3 variant with enhanced plasma membrane localization versus endosomal localization (Zhao et al., 2018). Given that polymorphisms in the genes encoding IFITM2 and IFITM3 have been proposed to increase their plasma membrane localization (Chesarino, McMichael et al., 2014a, Everitt, Clare et al., 2012, Jia, Xu et al., 2014, Wu, Grotefend et al., 2017), and that an additional polymorphism in the IFITM3 gene promoter results in significant loss of IFITM3 expression (Allen, Randolph et al., 2017), determining whether IFITMs affect SARS-CoV-2 cellular infections should be informative for understanding susceptibility and resistance of human cells to infection.

Published experimental results are contradictory with regard to the effect of IFITMs on SARS-CoV-2 cell entry. SARS-CoV-2 Spike-pseudotyped vesicular stomatitis virus infection was inhibited by human IFITM2 and IFITM3, but not IFITM1, in an overexpression screen of IFN effector proteins(Zang, Case et al., 2020). In contrast, IFITM3 overexpression did not inhibit infection of Spike-containing pseudovirus in a separate study (Zhao, Zheng et al., 2020). IFITMs have also been reported to inhibit cell-to-cell fusion induced by SARS-CoV-2 Spike protein(Buchrieser, Dufloo et al., 2020), though a conflicting report did not observe inhibition of Spike-mediated cell fusion when IFITM2 or IFITM3 were expressed in target cells (Zang et al., 2020). Overall, the impact of IFITMs on SARS-CoV-2 infections remains unclear and has yet to be examined with bona fide SARS-CoV-2. Herein, we measure the effects of mouse and human IFITMs on SARS-CoV-2 infections using both pseudovirus and genuine virus systems. We further utilize a series of mutants affecting critical functional amino acid motifs within IFITM3 and identify pro- and anti-viral effects that we clearly delineate in terms of distinct cellular locations and mechanisms of action.

## Results

### IFITMs inhibit SARS-CoV-2 Spike-pseudotyped lentivirus infection

To explore the effects of human IFITM1, IFITM2, and IFITM3 on SARS-CoV-2 infection, we transfected the SARS-CoV-2 receptor, ACE2, into HEK293T cells stably expressing FLAG-tagged IFITM constructs. We then infected the transfected cells with SARS-CoV-2 Spike-pseudotyped HIV-1 expressing luciferase (HIV-SARS-2), and measured luciferase production at 48 hpi. Compared to vector control cells, each of the IFITM-expressing lines showed significantly less luciferase production, indicating inhibition of Spike-mediated infection, with IFITM1 expression exhibiting particularly strong inhibition (Fig 1A). We also observed that all three of the IFITMs inhibited SARS-CoV Spike-pseudotyped lentivirus infection (HIV-SARS-1)(Fig 1B), with IFITM3 appearing to exhibit relatively more antiviral activity against HIV-SARS-1 than HIV-SARS-2 (Fig 1A,B). We confirmed uniform expression of each of the FLAG-tagged IFITM constructs by flow cytometry (Fig 1C). We further confirmed the ability of these IFITM constructs to limit HIV-SARS-2 infection in transiently-transfected Caco-2 cells, which naturally express ACE2 (Fig 1D). Furthermore, Caco-2 cells express basal IFITM3 protein at relatively high levels, while IFITM1 and IFITM2 are nearly absent. Knockdown of IFITM3 by siRNA transfection increased HIV-SARS-2 infection (Fig 1E). IFNβ treatment of Caco-2 cells strongly inhibited HIV-SARS-2 infection, which could be partially reversed with siRNA-mediated knockdown of IFITM3 (Fig 1E,F). These results indicate that basal, IFN-induced, and ectopically expressed IFITM3 inhibits HIV-SARS-2 infection of Caco-2 cells, and that, while multiple IFN stimulated genes are likely involved in repression of infection, IFITM3 is among those critical factors. In sum, our experiments indicate that IFITMs, both when overexpressed endogenously or ectopically, are restrictive of SARS-CoV-2 Spike-mediated virus infection

**Figure 1.**
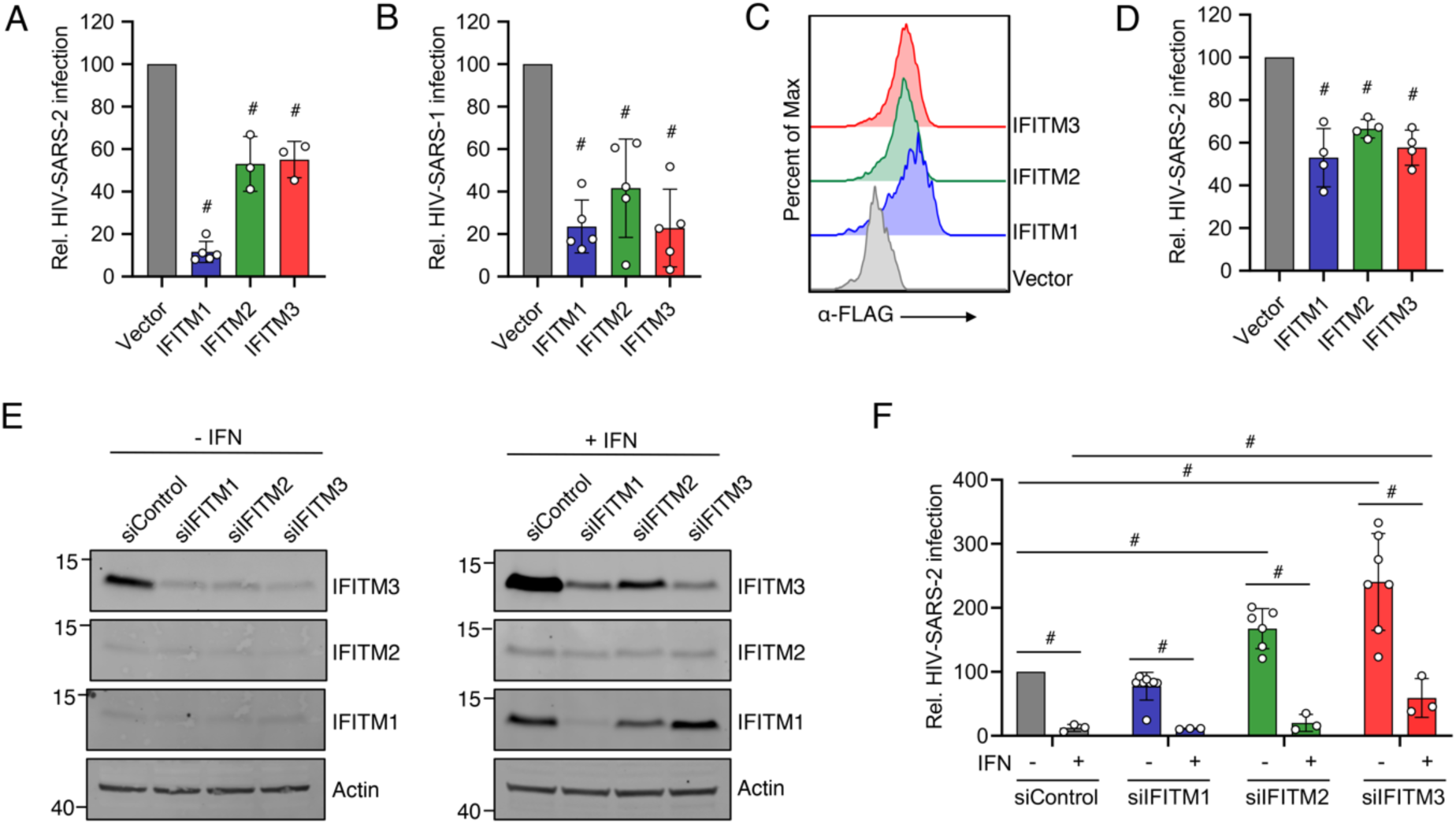
Human IFITMs inhibit SARS-CoV-2 Spike-pseudotyped lentivirus infections. HEK293T cells transduced with the indicated FLAG-tagged IFITM expression constructs or vector control plus transient transfection of ACE2 were infected with HIV-1expressing a luciferase reporter gene pseudotyped with **A)** SARS-CoV-2 Spike (HIV-SARS-2) or **B)** SARS-CoV Spike (HIV-SARS-1). Measurement of luciferase activity 72 h post infection served as a readout for infection. Luciferase readings were normalized to vector control, which was set to a value of 100. Bars in **A** and **B** represent averages with individual data points from 3-5 independent experiments shown as circles. Error bars represent SD. ^#^p<0.05 compared to vector control by ANOVA followed by Tukey’s multiple comparisons test. Not indicated in **A**, IFITM1 was also significantly different from IFITM2 and IFITM3. **C)** Representative example flow cytometry plots from cells transfected as in **A/B** demonstrate high transfection efficiency and expression of each of the FLAG-tagged IFITMs. **D)** Caco-2 cells were transfected as in **A/B** and infected with HIV-SARS-2. Luciferase readings at 72 h post infection were normalized to vector control, which was set to a value of 100. Bars represent averages with individual data points from 3 independent experiments shown as circles. Error bars represent SD. ^#^p<0.05 compared to vector control by ANOVA followed by Tukey’s multiple comparisons test. **E)** Caco-2 cells were transfected with the indicated siRNAs for 48 h followed by treatment with IFNβ for 24 h. Western blotting of cell lysates is shown. **F)** Cells as in **E** were infected with HIV-SARS-2. Luciferase readings at 48 h post infection were normalized to vector control, which was set to a value of 100. Bars represent averages with individual data points from 3-7 independent experiments shown as circles. Error bars represent SD. ^#^p<0.05 by ANOVA followed by Tukey’s multiple comparisons test for the comparisons indicated by lines.

### IFITMs inhibit SARS-CoV-2 infections

We next sought to validate results from pseudotyped virus infections using genuine SARS-CoV-2 at Biosafety Level 3. We utilized HEK293T cell lines stably expressing IFITMs (Fig 2A) that were transiently transfected with ACE2-GFP, and measured percent infection via staining for viral N protein and flow cytometry. Consistent with past reports suggesting low endogenous ACE2 expression in HEK293T cells, we found that infection was limited to cells that were ACE2-GFP positive (Fig 2B). We thus employed a gating strategy that focused on ACE2-GFP positive cells for measuring percent infection. Similar to results with Spike-pseudotyped lentivirus, we found that human IFITM1, IFITM2, and IFITM3 were each capable of inhibiting SARS-CoV-2 infections (Fig 2C,D). Additionally, we examined a cell line expressing a palmitoylation-deficient IFITM3 triple cysteine mutant (C71A, C72A, C105A; termed PalmΔ) that has been shown to lose activity against influenza virus and other viruses (Benfield, MacKenzie et al., 2020, Hach, McMichael et al., 2013, John, Chin et al., 2013, McMichael, Zhang et al., 2017, Percher, Ramakrishnan et al., 2016, Yount, Karssemeijer et al., 2012, Yount, Moltedo et al., 2010). Remarkably, IFITM3-PalmΔ maintained restriction of SARS-CoV-2 (Fig 2D). Given the highly unusual result with the IFITM3-PalmΔ cell line, we sought to authenticate the cell line by performing an influenza virus infection alongside one of our SARS-CoV-2 infections. Indeed, we confirmed the well-established results that IFITM3 strongly restricts influenza virus infection in a palmitoylation-dependent manner (Fig 2E) (Chesarino et al., 2017, Chesarino, McMichael et al., 2014b, Hach et al., 2013, McMichael et al., 2017, Melvin, McMichael et al., 2015, Percher et al., 2016, Yount et al., 2012, Yount et al., 2010). These results suggest that IFITMs provide palmitoylation-independent restriction of SARS-CoV-2 that is distinct from other viruses.

**Figure 2.**
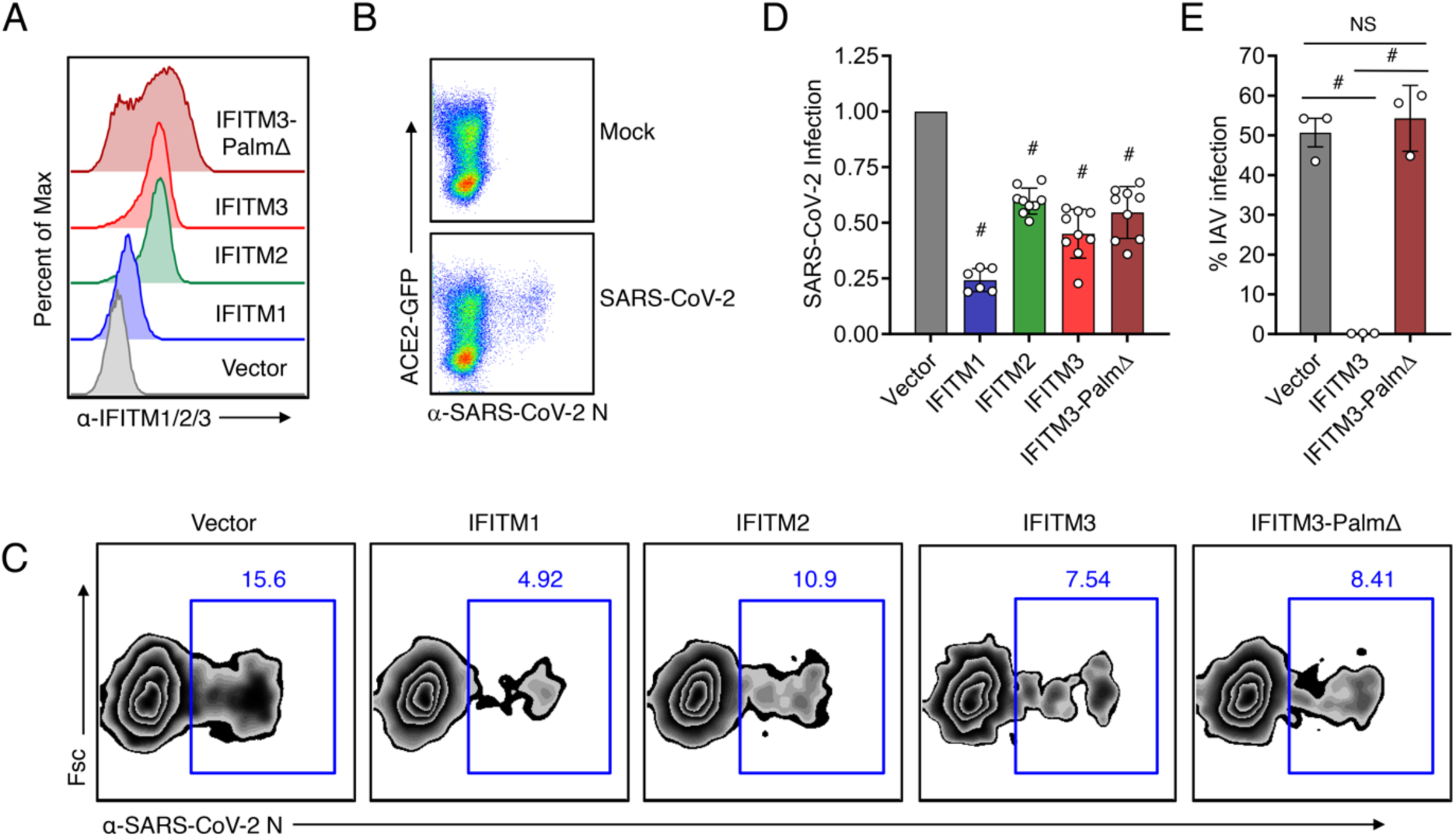
Human IFITMs inhibit infection by SARS-CoV-2. **A)** Flow cytometry histograms showing stable expression of the indicated IFITMs in HEK293T cells. **B)** Flow cytometry plots of HEK293T cells transfected with ACE2-GFP and infected with SARS-CoV-2 (MOI 0.5) for 24 h showing SARS-CoV-2 infection occurs in ACE2-GFP positive cells. **C)** Stable HEK293T lines as in **A** were infected with SARS-CoV-2 (MOI 1) for 24 h and analyzed for percent infection by flow cytometry. Representative example plots are shown for each cell line. **D)** Graph depicts normalized infection percentage measurements from 2-3 independent experiments, each performed in triplicate, from cells infected as in **C**. Bars represent averages with individual data points shown as circles. Error bars represent SD. ^#^p<0.05 compared to vector control by ANOVA followed by Tukey’s multiple comparisons test. **E)** The indicated cell lines were infected with influenza A virus (IAV) for 24h and percent infection was determined by flow cytometry. Graph depicts percent infection of triplicate samples. Bars represent averages with individual data points shown as circles. Error bars represent SD. ^#^p<0.05., NS, not significant, by ANOVA followed by Tukey’s multiple comparisons test.

### Human IFITM3 with mutations in its endocytic YxxΦ motif enhances SARS-CoV-2 infection

We additionally sought to confirm the effects of IFITMs on SARS-CoV-2 by transfecting IFITM constructs into stable HEK293T-ACE2-GFP cells. To measure SARS-CoV-2 infection, we employed a flow cytometry gating strategy in which cells that were expressing the highest levels of ACE2-GFP as well as IFITMs were analyzed for percent infection (Fig 3A,B). We again found that human IFITMs inhibit SARS-CoV-2 infection, though effects of transiently transfected IFITM2 did not reach statistical significance (Fig 3C,D). We also confirmed via transient transfection that IFITM3-PalmΔ is active in restricting the virus (Fig 3C,D). We further found that, consistent with past results examining endogenous ACE2 (Huang et al., 2011), IFITM expression did not affect levels of exogenously expressed ACE2-GFP (**Fig E,F**). We additionally examined mouse Ifitm1-3 and found that restriction of SARS-CoV-2 infection is a conserved activity of mouse and human IFITMs, and that restriction by mouse IFITM3 occurred independently of palmitoylation like its human counterpart (Fig 4A-C).

**Figure 3.**
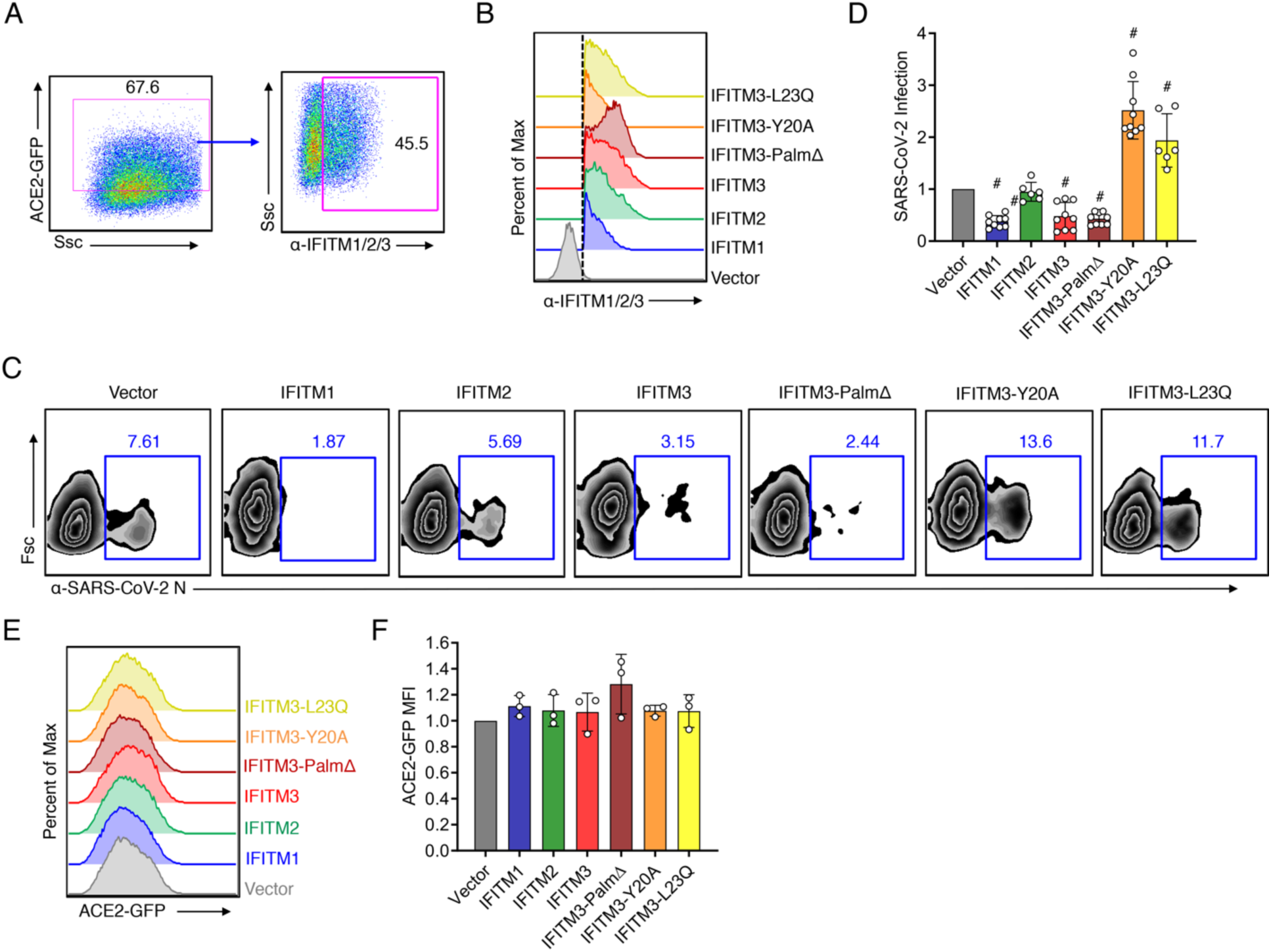
Mutation of the human IFITM3 endocytosis motif converts IFITM3 into an enhancer of SARS-CoV-2 infections. HEK293T-ACE2-GFP stable cells were transiently transfected with human IFITM plasmids or vector control for 24 h. **A)** Flow cytometry gating strategy for analyzing cells with high ACE2-GFP expression and IFITM expression. **B)** Flow cytometry histograms showing expression of each of the IFITM constructs, and the gating as indicated in **A** and by the dotted line, which was used for examining percent infections in IFITM-expressing cells. **C)** Transfected HEK293T-ACE2-GFP cells were infected with SARS-CoV-2 (MOI 1) for 24h and analyzed for percent infection by flow cytometry. Representative example plots are shown for each transfection condition. **D)** Graph depicts normalized infection percentage measurements from 2-3 independent experiments, each performed in triplicate, from cells infected as in **C**. Bars represent averages with individual data points shown as circles. Error bars represent SD. ^#^p<0.05 compared to vector control by ANOVA followed by Tukey’s multiple comparisons test. **E)** Representative example flow cytometry histograms of ACE2-GFP from non-infected samples. **F)** Normalized measurements of ACE2-GFP mean fluorescence intensity (MFI) of non-infected samples from three separate experiments. Bars represent averages with individual data points shown as circles. Error bars represent SD. Statistical significance (p<0.5 by ANOVA followed by Tukey’s multiple comparisons test) was not observed for comparisons between any samples.

**Figure 4.**
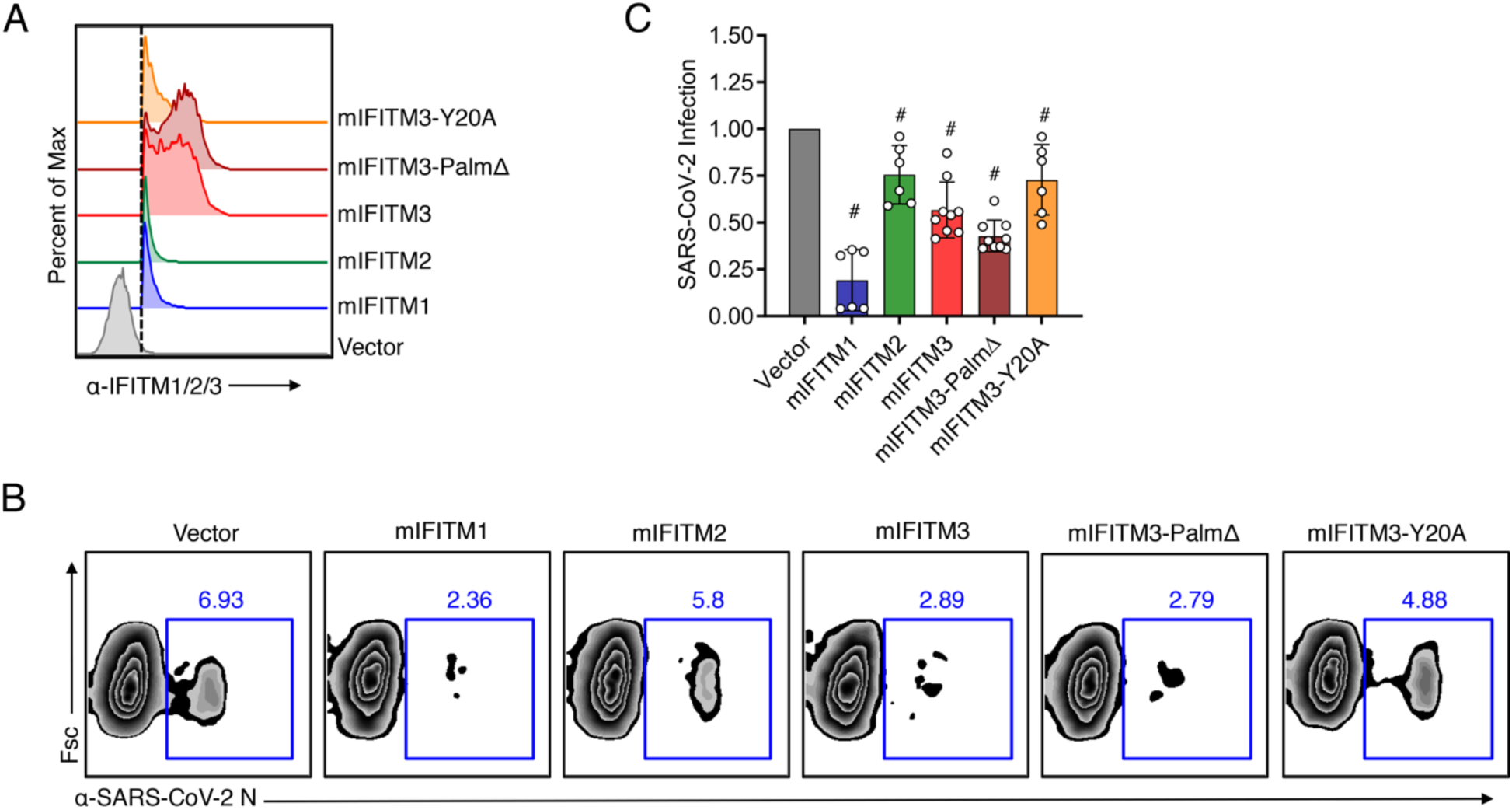
Mouse IFITMs inhibit SARS-CoV-2 infection. HEK293T-ACE2-GFP stable cells were transiently transfected with mouse (m) IFITM plasmids or vector control for 24 h. **A)** Flow cytometry histograms showing expression of each of the IFITM constructs with the dotted line indicating gating that was used for examining percent infections in IFITM-expressing cells. **B)** Transfected HEK293T-ACE2-GFP cells were infected with SARS-CoV-2 (MOI 1) for 24h and analyzed for percent infection by flow cytometry. Representative example plots are shown for each transfection condition. **C)** Graph depicts normalized infection percentage measurements from 2-3 independent experiments, each performed in triplicate, from cells infected as in **B**. Bars represent averages with individual data points shown as circles. Error bars represent SD. ^#^p<0.05 compared to vector control by ANOVA followed by Tukey’s multiple comparisons test.

We next examined mouse and human IFITM3 mutants that impact protein localization. Y20 is a critical amino acid within a YxxΦ endocytosis motif (20-YEML-23 in human IFITM3), and has been shown in several studies to be required for active endocytosis of IFITM3(Chesarino et al., 2014a, Jia, Pan et al., 2012, Jia et al., 2014, McMichael, Zhang et al., 2018). Mutation of Y20 or blocking this residue via phosphorylation by Src kinases results in accumulation of IFITM3 at the plasma membrane (Chesarino et al., 2014a, Compton, Roy et al., 2016, Jia et al., 2012, Jia et al., 2014, McMichael et al., 2018). Human IFITM3-Y20A caused an increase in SARS-CoV-2 infection as compared to vector control cells, suggesting that IFITM3 at the plasma membrane promotes, rather than inhibits, infection (Fig 3C,D). In contrast, mouse IFITM3-Y20A did not increase infection compared to vector control cells (Fig 4B,C), revealing a species-specific distinction between mouse and human IFITM3 in their ability to promote SARS-CoV-2 infection. To confirm that enhancement of infection was due to disruption of the 20-YEML-23 endocytosis motif, we tested an additional construct, human IFITM3-L23Q, which similarly accumulates at the plasma membrane due to disruption of the critical Φ residue of the endocytosis signal (Chesarino et al., 2014a). IFITM3-L23Q expression also increased SARS-CoV-2 infection, confirming the unusual ability of IFITM3 to enhance infection of this virus (Fig 3C,D). Overall, these results demonstrate that while mouse and human IFITMs generally restrict infection, human IFITM3 can promote infection when its localization is shifted toward the plasma membrane.

### Human IFITM3 with mutations in its endocytic YxxΦ motif enhances SARS-CoV-2 Spike-mediated cell-to-cell fusion

We next examined whether IFITM3 affects SARS-CoV-2 Spike-mediated membrane fusion in a cell-to-cell fusion assay. U2OS cells co-transfected with Spike and mCherry expression plasmids were mixed with Calu-3 target cells that were co-transfected with plasmids encoding EGFP and IFITMs. After 24 h co-culture, mCherry and EGFP double positive multi-nucleated cell syncytia could be observed dependent upon the presence of Spike protein, demonstrating Spike-mediated fusion between the two cell types (Fig 5A). Upon transfection of IFITM3 into Calu-3 target cells, double positive syncytia remained readily observable upon visual inspection, though quantification of nuclei number within these double positive syncytia suggested that IFITM3 may have a modest ability to limit S-mediated cell-to-cell fusion (Fig 5A,B). In contrast, transfection of IFITM3-Y20A or IFITM3-L23Q resulted in an increase in the size of double positive syncytia as compared to vector control or WT IFITM3 transfections, which was confirmed by quantification of nuclei per syncytia (Fig 5A,B). These results provide confirmation of infection experiments indicating that IFITM3 enriched at the plasma membrane due to endocytosis-impairing mutations enhances SARS-CoV-2 Spike-mediated membrane fusion.

**Figure 5.**
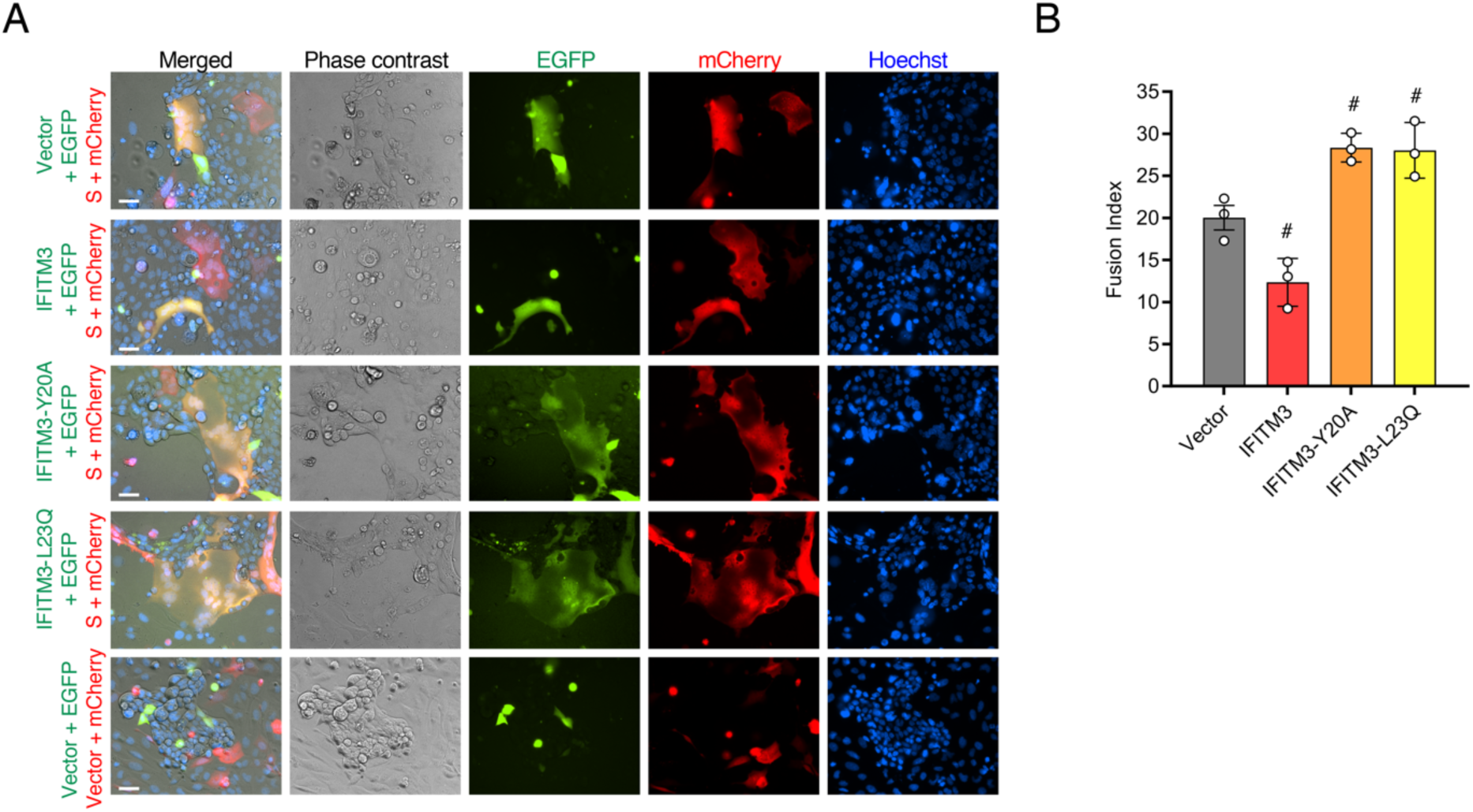
Mutation of the human IFITM3 endocytosis motif converts IFITM3 into an enhancer of SARS-CoV-2 Spike-mediated cell-to-cell membrane fusion. U2OS cells transfected with EGFP and the indicated IFITM3 constructs or vector control were mixed Calu-3 cells transfected with mCherry and SARS-CoV-2 Spike (S) or vector control. Cells were co-cultured for 24 h and imaged by fluorescent microscopy to visualize EGFP and mCherry as well as nuclei via Hoechst dye. **A)** Representative example fluorescent imaging of the indicated co-cultures showing the presence of EGFP/mCherry double positive cell syncytia dependent upon S expression as well as varying effects of the IFITM3 constructs. Scale bar indicates 50 μm. **B)** Quantification of nuclei per EGFP/mCherry double positive cell syncytia in co-cultures involving the indicated IFITM3 constructs as compared to vector control. Bars depict averages of three independent experiments with circles representing the individual data points. For each condition, nuclei within 50-75 individual syncytia were counted. Error bars represent SD. ^#^p<0.05 compared to vector control by ANOVA followed by Tukey’s multiple comparisons test.

### Anti-SARS-CoV-2 activity of IFITM3 requires its amphipathic helix domain and is counteracted by TMPRSS2 overexpression

Our results suggest that IFITM3 is able to inhibit SARS-CoV-2 infection overall, but that shifting human IFITM3 localization away from the endosomal system and toward the plasma membrane results in enhancement of infection (Fig 2, 3). Given that SARS-CoV-2 has dual cellular entry pathways involving Spike proteolytic activation either at the plasma membrane or within endosomes (Hoffmann, Kleine-Weber et al., 2020, Shang, Wan et al., 2020), these findings may indicate that IFITM3 has distinct effects on the virus dependent upon the location at which virus fuses with the cell membrane. Plasma membrane-associated processing of Spike involves TMPRSS2 protease, and it has been reported that TMPRSS2 overexpression increases the proportion of virus able to fuse at the cell surface (Hoffmann et al., 2020). Further, it is reported that high TMPRSS2 levels allow coronaviruses to avoid restriction by IFITMs (Bertram et al., 2013, Zheng et al., 2020). In line with these data, we found that TMPRSS2 overexpression decreased the ability of IFITM3 to inhibit SARS-CoV-2 infection (Fig 6A,B). Interestingly, IFITM3 slightly enhanced infection in the presence of TMPRSS2 overexpression in 2 out of 3 experiments (Fig 6B). Inhibition of infection by IFITM3-PalmΔ and IFITM1 was also counteracted by TMPRSS2, though enhancement of infection was not observed for these constructs (Fig 6A,B).

**Figure 6.**
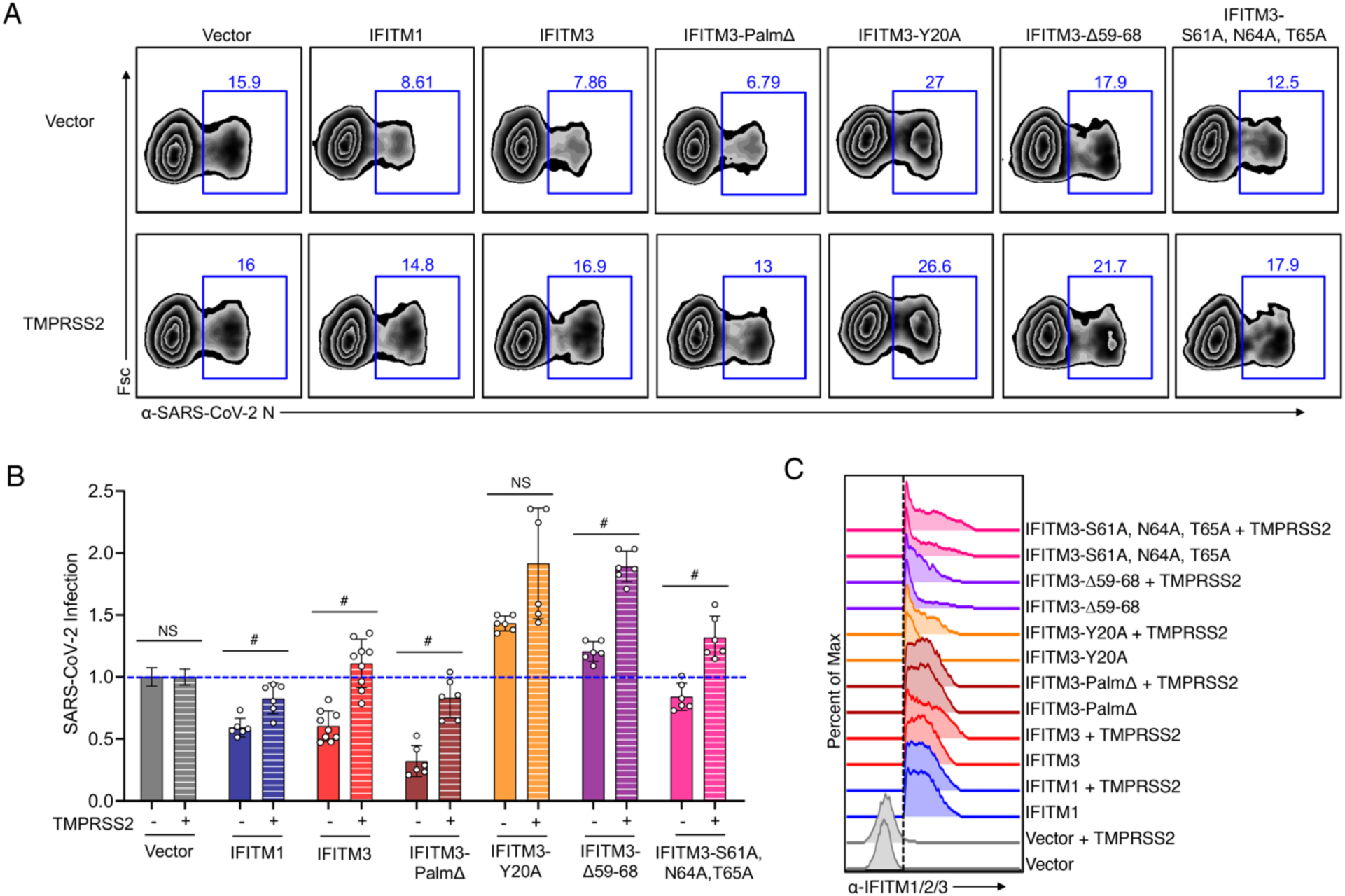
IFITM3 restriction of SARS-CoV-2 infection requires the IFITM3 amphipathic helix while enhancement of infection does not. HEK293T-ACE2-GFP stable cells were transiently transfected with human IFITM plasmids or vector control plus TMPRSS2 or vector control for 24 h. **A)** Transfected HEK293T-ACE2-GFP cells were infected with SARS-CoV-2 (MOI 1) for 24h and analyzed for percent infection by flow cytometry. Representative example plots are shown for each transfection condition. **B)** Graph depicts normalized infection percentage measurements from 2-3 independent experiments, each performed in triplicate, from cells infected as in **B**. Bars represent averages with individual data points shown as circles. Error bars represent SD. The blue dotted line is added to the graph to assist with comparisons to vector control levels of infection. ^#^p<0.05 by ANOVA followed by Tukey’s multiple comparisons test. NS, not significant. Not indicated, additional comparisons of interest with p<0.05 include IFITM3 (-) vs. IFITM3-Y20 (-), IFITM3Δ59-68 (-), and IFITM3-S61A,N64A,T65A (-). **C)** Flow cytometry histograms showing expression of each of the IFITM constructs with the dotted line indicating gating that was used for examining percent infections in IFITM-expressing cells.

The IFITM3 amphipathic helix domain located at amino acid positions 59-68 is required for inhibition of influenza virus infection (Chesarino et al., 2017). This helix was recently shown to be able to alter mechanical properties of lipid membranes, such as curvature and lipid order, thus inhibiting viral membrane fusion(Guo et al., 2020). In the context of SARS-CoV-2, we found that expression of an amphipathic helix-deleted mutant of IFITM3 (Δ59-68) resulted in a modest enhancement rather than inhibition of infection (Fig 6A,B). Likewise, decreasing amphipathicity of the helix by mutation of its three hydrophilic residues (S61A, N64A, T65A), decreased restriction of infection (Fig 6A,B), further demonstrating that the amphipathicity of IFITM3, and likely its associated membrane-altering properties, underlie restriction of SARS-CoV-2. Notably, both of the amphipathic helix mutants were increased in their ability to enhance SARS-CoV-2 infection when co-expressed with TMPRSS2 (Fig 6A,B), providing the first clear evidence that IFITM3 enhancement of CoV infection occurs via a mechanism distinct from its amphipathicity-based membrane alterations. These results support a model wherein IFITM3 exerts opposing activities on SARS-CoV-2, including amphipathicity-dependent restriction of virus at endosomes and amphipathicity-independent enhancement of infection at the plasma membrane.

## Discussion

In the present study we have identified IFITM1, IFITM2, and IFITM3 as restrictors of SARS-CoV-2 infection of cells. Our result showing that IFITM3 inhibits SARS-CoV-2 infection independently of *S*-palmitoylation is surprising since intact palmitoylation sites are necessary for inhibition of a number of viruses, including influenza virus, dengue virus, CoV-NL63, and CoV-229E (Benfield et al., 2020, Hach et al., 2013, John et al., 2013, McMichael et al., 2017, Percher et al., 2016, Yount et al., 2012, Yount et al., 2010, Zhao et al., 2018). This conclusion is supported by experiments with three distinct expression systems (stable transduction of human IFITM3-PalmΔ in the pLenti vector and transient transfections of human and mouse IFITM3-PalmΔ in pCMV) (Fig 2,3,4). Previous work suggests that S-palmitoylation increases the rate at which IFITM3 traffics to influenza virus-containing endosomes during infection (Spence, He et al., 2019). Our results suggest possible differences between influenza virus and SARS-CoV-2 either in their endocytic entry pathways or fusion kinetics, and IFITM constructs may prove useful in dissecting such differences.

Mechanistically, we found that the IFITM3 amphipathic helix is necessary for its SARS-CoV-2 restrictive capacity (Fig 6). We previously showed that this helix was also required for restriction of influenza virus and proposed that the helix affected virus fusion by wedging into the inner leaflet of the membrane bilayer, thus generating local membrane curvature that is disfavorable to virus-to-cell fusion (Chesarino et al., 2017). More recent research has shown that the amphipathic helix is indeed able to mechanically induce membrane curvature and increase lipid order (Guo et al., 2020). Overall, IFITMs likely restrict SARS-CoV-2 infection by mechanically disfavoring viral membrane fusion reactions occurring at endosomes.

Remarkably, we observed that IFITM3-Y20A and -L23Q constructs enhance SARS-CoV-2 infection (Fig 4). Since these constructs are well characterized to accumulate at the plasma membrane of cells due to loss of interaction with the AP-2 endocytic adaptor protein complex (Chesarino et al., 2014a, Jia et al., 2014, McMichael et al., 2018), these data suggest that IFITM3 localization to the plasma membrane enhances infection. Given that WT IFITM3 traffics to the plasma membrane prior to its endocytosis, and since phosphorylation of IFITM3 Y20 by Src kinases also blocks endocytosis (Chesarino et al., 2014a, Jia et al., 2012), there are likely opposing restrictive and enhancing activities occurring in cells expressing WT IFITM3. It will be interesting to determine whether these balances are shifted in different cell types with distinct endocytic or phosphorylation capacities.

Enhancement of SARS-CoV-2 infection at the plasma membrane does not require the IFITM3 amphipathic helix, as demonstrated by our observation that TMPRSS2 overexpression converts amphipathic helix mutants into strong enhancers of infection (Fig 6). We did not observe an enhancing effect of human IFITM1, which also has a conserved and functional amphipathic helix domain (Chesarino et al., 2017), further supporting that enhancement of infection at the plasma membrane is not occurring via amphipathic helix-mediated mechanical membrane alterations (Fig 6). These data are in alignment with past reports that IFITM3, but not IFITM1, enhances CoV-OC43 infections (Zhao et al., 2014). Removal of the IFITM1 extended C-terminal tail converted IFITM1 into an enhancer of specific CoV infections (Zhao et al., 2014, Zhao et al., 2018). Likewise, and similar to our results with SARS-CoV-2, mutation of human IFITM3 Y20 was also reported to enhance SARS-CoV and MERS-CoV pseudotype virus infections, but not CoV-229E or CoV-NL63 pseudotype infections (Zhao et al., 2018). Since enhancement of infection does not correlate with specific receptor usage of the different coronaviruses, it is unlikely that IFITMs directly affect the virus receptors. In that regard, we did not observe an effect of IFITM proteins on levels of exogenously expressed ACE2-GFP (Fig 3E,F). It may be possible that the short IFITM3 C-terminal tail, predicted to be exposed at the cell surface, directly interacts with specific features of CoV surface glycoproteins or with entry factors. It is also interesting to note that mouse IFITM3 has an extended C-terminal tail compared to human IFITM3, consistent with results that mouse IFITM3-Y20A did not enhance infection (Fig 4B,C). Overall, our results indicate that IFITMs are not likely exerting a broad mechanism of amphipathicity-based membrane alteration to promote infection, but rather are more specifically coopted by specific CoVs, such as SARS-CoV-2, under certain conditions. An early study on a limited cohort suggests that the human *IFITM3* rs12552-C SNP, which has been proposed to result in IFITM3 plasma membrane localization (Chesarino et al., 2014a, Everitt et al., 2012, Jia et al., 2012), is a risk factor for severe COVID-19(Zhang, Qin et al., 2020). Larger studies are warranted to examine whether this and other polymorphisms in *IFITM3* (Zani & Yount, 2018) positively or negatively influence COVID-19 severity.

## Materials and Methods

### Biosafety

All work with live SARS-CoV-2 was performed at Biosafety Level 3 (BSL3) according to standard operating procedures approved by the Ohio State University BSL3 Operations Group and Institutional Biosafety Committee. Samples were removed from the BSL3 facility for flow cytometry analysis only after decontamination with 4% paraformaldehyde for a minimum of 1 h according to an in-house validated and approved method of sample decontamination.

### Cell lines and cell culture

All cells were cultured at 37°C with 5% CO2 in a humidified incubator and were grown in Dulbecco’s Modified Eagle’s Medium (DMEM) supplemented with 10% FBS (Atlas Biologicals). HEK293T, Calu-3, and Caco-2 were purchased from ATCC. The origin of our U2OS cells are unknown, but the identity and purity of the cells were verified by STR profiling. Vero E6 cells were a kind gift from Dr. Mark Peeples (Nationwide Children’s Hospital) and were treated with Mycoplasma Removal Agent (MP Biomedical) for 1 month to ensure lack of contamination before virus propagation. All other cell lines were routinely treated with Mycoplasma Removal Agent for 1 week upon thawing of new cultures. HEK293T cells stably expressing human IFITM1, IFITM2, IFITM3, IFITM3-PalmΔ, or empty pLenti-puro vector that were used in genuine virus infections were generated and validated as described previously (McMichael et al., 2018), with the exception that the IFITM3-PalmΔ cell line required three sequential transductions to achieve expression comparable to WT IFITM3 in the majority of the cells. HEK293T cells stably expressing GFP-tagged human ACE2 (Origene) were generated using the same methodology. HEK293T cell lines stably transduced with FLAG-tagged human IFITM1, IFITM2, IFITM3, or pQCXIP vector control for HIV-SARS-1 and HIV-SARS-2 infection experiments were described previously (Shi, Ozog et al., 2018). Type-I interferon (human recombinant interferon beta, NR-3085, BEI Resources) was added to Caco-2 cells at 48h post siRNA transfection at a concentration of 250 IU/mL. Cells were incubated with interferon for a 24h period prior to being removed and replaced with virus inoculum.

### SARS-CoV-2 propagation and titering

SARS-CoV-2 USA-WA1/2020 stock from BEI Resources was diluted 1:10,000 in DMEM and added to confluent Vero cells for 1 h at 37°C. Following the 1 h infection period, virus-containing media was replaced with DMEM containing 4% FBS and incubated at 37°C for 72 h at which point significant cytopathic effect was observed. Cell supernatant containing infectious virus was centrifuged at 1,000 x g for 10 min to remove cell debris, and was aliquoted, flash frozen in liquid nitrogen, and stored at -80°C. Virus titer was determined by TCID50 assay in Vero cells using the presence of cytopathic effect coupled with viral N protein staining for identification of infected wells. The viral stock titer was further confirmed by standard plaque assay with 0.3% agarose (Sigma) overlay and visualization with 0.25% Crystal Violet (Sigma) in Vero cells. Full genome sequencing of the virus stock by the Ohio State University Medical Center Clinical Diagnostics Laboratory confirmed an absence of major mutations or deletions, including within the Spike protein furin cleavage site.

### Plasmids, transfections, and siRNA knockdowns

Plasmid encoding untagged ACE2 was a gift from Hyeryun Choe (Addgene plasmid # 1786; http://n2t.net/addgene:1786;RRID:Addgene_786). Transfection of HEK293T cell lines stably expressing human IFITM1, IFITM2 or IFITM3 (35) with ACE2 was performed with TransIT-293 reagent (Mirus). Transfection of Caco-2 cells with empty pQCXIP or pQCXIP-IFITM1, -IFITM2, -IFITM3 (gifts from Chen Liang) was performed using Lipofectamine 2000 (Thermo Fisher). For SARS-CoV-2 infection experiments, pCMV expression plasmid for N-terminally HA-tagged human and mouse IFITM1, IFITM2, IFITM3, IFITM3-PalmΔ and IFITM3-Y20A as well as human IFITM3-L23Q, IFITM3-Δ59-68, and IFITM3-S61A,N64A,T65A were previously described (Chesarino et al., 2017, Chesarino et al., 2014a, Hach et al., 2013, Percher et al., 2016, Yount et al., 2010). TMPRSS2 in pCAGGS was a kind gift from Dr. Stephan Pöhlmann (Georg-August-Universität Göttingen). ACE2-GFP in pLenti-puro was purchased from Origene. All transient transfections for SARS-CoV-2 infection experiments were performed using LipoJet transfection reagent (Signagen Laboratories). siRNAs targeting *IFITM1* (s16192), *IFITM2* (s20771), *IFITM3* (s195035) and a non-targeting siRNA control pool (4390844) were purchased from Ambion. siRNA-mediated knockdown was performed by transfecting Caco-2 cells with 20 nM amounts of siRNA and Lipofectamine RNAiMAX (Thermo Fisher). Knockdown efficiency was assessed by Western blot analysis at 72h post transfection.

### Pseudotyped virus infections and luciferase assays

Virus-like particles pseudotyped with SARS-CoV-2 or SARS-CoV-1 Spike protein were produced by transfecting HEK293T cells with a plasmid encoding HIV-1 NL4-3 lacking *vpr* and encoding luciferase in place of *env* (a gift from Vineet KewalRamani) and pcDNA-SARS-CoV2-C9 or pcDNA-SARS-CoV1-C9 (gifts from Thomas Gallagher) using TransIT-293 reagent. Cells were incubated with virus-containing medium for 72h, after which cells were lysed with Passive Lysis Buffer (Promega). Luciferase assays were performed using Luciferase Assay System (Promega). Luciferase values were measured with a Tecan Infinite M1000 plate reader.

### SARS-CoV-2 infections and flow cytometry

Cells were infected with SARS-CoV-2 for 24 hours, after which cells were fixed with 4% paraformaldehyde (Thermo Scientific) for 1 h at room temperature, permeabilized with PBS containing 0.1% TritonX-100, and blocked with PBS/2% FBS. Cells were then stained with mouse-produced anti-SARS-CoV-2 N (Sino Biological) and a cocktail of rabbit-produced anti-IFITM2/3 (ProteinTech) and anti-IFITM1 (Cell Signaling Technologies). Primary antibody labeling was followed by labeling with anti-mouse-AlexaFluor-647 and anti-rabbit-AlexaFluor-555 secondary antibodies (Life Technologies). Flow cytometry data was collected using a FACSCanto II flow cytometer (BD Biosciences). Data analysis was performed using FlowJo software. Infected cell gates were set using non-infected control samples.

### Cell fusion assay

Using Lipofectamine 3000 (Life Technologies), U2OS cells were co-transfected with pmCherry and either empty pCAGGS vector or SARS-CoV-2 Spike Glycoprotein (BEI Resources, deposited by Dr. Florian Krammer, Icahn School of Medicine at Mount Sinai); Calu-3 cells were co-transfected with pEGFP and either empty pCMV vector, IFITM3, IFITM3-Y20A, or IFITM3-L23Q. Transfected Calu-3 and U2OS cells were mixed and co-incubated for 24 hours. Following incubation, the co-cultures were contra-stained with Hoechst 33342 and imaged using a Nikon Eclipse Ti-E inverted microscope (Nikon Instruments). Nikon NIS Elements AR software was used to quantify fusion indices calculated as number of nuclei per double-positive (expressing both EGFP and mCherry) cell fusion cluster.

## Acknowledgements

We would like to thank Peter Lai and Alan Rein for their assistance in producing lentiviral pseudotypes containing SARS-CoV-1 and SARS-CoV-2 Spike proteins. We thank Dr. Eugene Oltz for critical reading of the manuscript.

